# Lifestyle and biological factors influence the relationship between mental health and low-grade inflammation

**DOI:** 10.1101/609768

**Authors:** A Gialluisi, M Bonaccio, A Di Castelnuovo, S Costanzo, A De Curtis, M Sarchiapone, C Cerletti, MB Donati, G de Gaetano, L Iacoviello, on behalf of the Moli-Sani Study Investigators

## Abstract

Mental health modulates the risk of common chronic conditions like cardiovascular disease, cancer and diabetes. Although inflammation is thought to partly explain this link, its relation with mental health is still unclear and largely unexplored.

We investigated three scales assessing psychological resilience (CD-RISC), depression symptoms (PHQ8) and mental wellbeing (SF36-MCS) in an Italian adult population cohort (N_max_=16,952). We performed stepwise generalized linear models to test the association between each scale and INFLA-score, a composite blood-based inflammation index. At each step, a class of potential mediators was included in the model, namely health conditions, lifestyle factors, or both (full model). Full model analysis was also conducted on single blood markers involved in the inflammatory process.

In the baseline model, we observed significant associations of PHQ8 (standardized β=0.024, p=8.9×10^−3^) and SF36-MCS (β = −0.021, p=7×10^−3^) with INFLA-score. These associations survived adjustment for health conditions but not for lifestyle factors, which explained 81% and 17% of the association with PHQ8 and SF36-MCS, respectively. Significant associations (p<4.2×10^−3^) after mediator adjustment were observed for single low-grade inflammation markers, including platelet distribution width (with PHQ8 and CD-RISC), granulocyte-and neutrophil-to-lymphocyte ratios, monocyte and lymphocyte fractions (with SF36-MCS).

These findings suggest that the relationship between mental health and low-grade inflammation is largely influenced by lifestyle. However, the associations with specific biomarkers related to inflammation are partly independent and might be explained by biological factors. Interestingly, these associations are in line with recent blood transcriptomic analyses of depressed subjects, reporting up- and down-regulation of genes related to innate and adaptive immunity, respectively.

## Introduction

Psychological health represents an important component of the general health status. Indeed, according to the World Health Organization, “health is a state of complete physical, mental and social wellbeing, and not merely the absence of disease or infirmity”. In the last decades, an increasing amount of research has focused on the link between frequent mental disorders, like anxiety and depression^1^, and common health conditions such as cardiovascular diseases (CVD)^2^. Depression (or Major Depressive Disorder, MDD) probably represents the most investigated mental disease from this point of view, showing a high comorbidity with chronic conditions like cardiovascular disease (CVD)^3^, cancer^4^ and diabetes^5^. Although this link may look obvious, given the prognosis of these conditions, longitudinal studies report an increased incidence of these disorders for depressed people, suggesting that MDD is a risk factor for and may share common biological pathways with these conditions. Among these pathways, low-grade inflammation has been proposed as one of the most plausible links^3–6^. Several clinical and preclinical studies have associated high levels of circulating pro-inflammatory cytokines like interleukin-1β (IL-1β), interleukin-6 (IL-6) and tumor necrosis factor-α(TNF-α), with depressive symptoms and the efficacy of antidepressant treatments in humans, as well as with depression-like behaviours in animal models^6–8^. Similarly, circulating levels of C-reactive protein (CRP) and measures of blood platelet indices, such as mean platelet volume (MPV), are reportedly increased in depressed subjects^6,9–11^. Other markers of low-grade inflammation were associated with depression, including neutrophil-to-lymphocyte ratio (NLR) and red cell distribution width (RDW)^10,12^. However, other studies have reported inconsistent results and many open questions remain in this field^6–8^.

Other scales closely related to mental health, like health-related quality of life (QoL, a measure of mental and physical wellbeing) have been found to be associated with inflammation, both in the general population^13^, and in selected samples, including elders^14^, and people affected by diabetes^15^, metabolic syndrome^16^, depression^17^ and schizophrenia^18^. These studies, which investigated mainly inflammatory markers like CRP, reported association with both mental and physical components of QoL^15,18,19^, although not always consistently. As an example, Faugere et al.^17^ detected a significant association between QoL and CRP levels in a sample of depressed subjects, but only with the physical component. More in general, only one study investigated the relation between wellbeing and inflammation in a general population setting, reporting a positive association between self-rated life satisfaction, CRP and fibrinogen levels in the Scottish population^13^.

Similarly, psychological resilience – which represents the ability to cope with continued and/or intense exposure to environmental stressors - has been closely connected to inflammation^20,21^, mainly based on findings in animal models (reviewed in^22^). A few studies in humans supported this association, either directly, by reporting a moderation effect of resilience on the association between adverse childhood experiences and IL-6 circulating levels^23^, or indirectly, by revealing an association with transcriptional down-regulation of pro-inflammatory genes, and with up-regulation of innate antiviral and antibody-related genes^24^. However, other studies found contrasting results, e.g. positive correlations of resilience with the circulating levels of pro-inflammatory cytokines like interferon γ(IFN-γ) and IL-6^25^. Of note, we found no study which analysed the link between resilience and inflammation in a general population setting.

Overall, much remains to be done in order to clarify the relation between mental health and low-grade inflammation. Among the open issues, the most important remain the direction of associations, the role of potential confounders and mediators in this relation, and the paucity of studies in the general population and of inflammation biomarkers tested, especially with reference to resilience and wellbeing. This limits our ability to disentangle the relation of mental health with the different components of inflammation.

Here, we analysed this relation in a large population-based cohort of adult Italians. We tested the association between subclinical inflammation and three measures of mental health, namely i) psychological resilience^26^; ii) depressive symptoms^27^; and iii) mental quality of life^28^. We analysed the global burden of potential mediators on this association, including both prevalent health conditions and main lifestyle factors. Then, to increase the resolution of our analysis, we tested associations of these three scales with a number of circulating blood markers involved in inflammatory processes to different degrees. Our findings allow to progress towards the comprehension of the complex relation between mental health and low-grade inflammation.

## Methods and Materials

### Study Population

The subjects under investigation were enrolled in the Moli-sani Study, a population-based cohort which randomly included 24,325 subjects living in Molise, a small region located in central Italy, with about 300,000 citizens. Between March 2005 and April 2010, men and women aged ≥ 35 years were randomly recruited from city-hall registries, using electronically generated numbers, with an overall response rate of 70%. Exclusion criteria were pregnancy, disturbances in understanding/willing processes, ongoing poly traumas or coma^29,30^. The Moli-sani Study was approved by the Catholic University ethical committee and all the participants provided written informed consent.

For the purpose of this study, we selected subjects for whom complete questionnaire data on psychological resilience (N=11,272), depression symptoms (N=13,776) or health-related mental quality of life (N=19,035) were available (see paragraph *Psychometric scores* for further details).

Subjects with missing information for assessments on dietary patterns, physical activity or smoking habits and missing values of anthropometric measures such as body mass index (BMI) and waist-to-hip ratio were filtered out, as well as those individuals with missing data for at least one of the low-grade inflammation biomarkers tested here (see paragraph below). Finally, in order to investigate the relation between low-grade inflammation and mental health in an unbiased way, we filtered out before analysis samples showing signs of acute ongoing inflammation (C-reactive protein levels > 10 mg/L). After these filters, 10,415 samples were available for the analysis of resilience, 12,732 for depressive symptoms and 16,952 for mental quality of life.

### Blood measures and low-grade inflammation index

To model the relation between each of the psychometric scales analysed and low-grade inflammation, we used the INFLA-score - a global index of low-grade inflammation based on four circulating biomarkers, namely C-reactive protein levels (CRP), blood platelet count (Plt), white blood cell count (WBC) and granulocyte-to-lymphocyte ratio (GLR) - capturing both plasma and cellular circulating inflammation^30^. This score consists of an equal weight combination of these four biomarkers, as briefly explained below. The 10-tiles of all circulating levels of the four biomarkers of interest were computed. For each of the four components of INFLA-score, the distribution of biomarker levels was divided into 10-tiles and each 10-tile was assigned a corresponding score, as follows: samples with a biomarker level laying in Q5 and Q6 (median values) were labelled with score 0, biomarker levels >Q6 (higher tail of distribution) were assigned an increasing score from +1 (Q7) through +4 (Q10), while biomarker levels <Q5 (lower tail of distribution) were assigned a negative score from −4 (Q1) to −1 (Q4). Then the INFLA-score was computed as a sum of the resulting scores over the four biomarkers tested, with total values ranging between −16 (lowest grade of subclinical inflammation) and +16 (highest grade of subclinical inflammation).

In addition to the INFLA-score and its component biomarkers, other blood markers which are involved to different extents in inflammatory processes were used in specific analyses (see below), to have a more fine-grained picture of the relation between inflammation and mental health. These additional parameters included mean platelet volume (MPV), plateletcrit (PCT), platelet distribution width (PDW), red cells distribution width (RDW), fraction of lymphocytes (LY), monocytes (MO), granulocytes (GR) and neutrophils (NE) over the total WBC, and neutrophil-to-lymphocyte ratio (NLR), and their selection was motivated by the well-recognized role of platelets in inflammatory processes^31,32^, and by the strong link between inflammation and immunity^33,34^. Further details on blood measurements are reported in *Supplementary Methods* and in^35^.

### Psychometric scores

Three main self-administered psychometric tests were used as exposure variables, representing distinct but related psychological domains, namely psychological resilience, depressive symptoms and mental health.

Psychological resilience was tested through the Connor-Davidson Resilience Scale (CD-RISC), a self-rated assessment based on 25 items. Since each item is rated on a 5-point scale (0–4), the total score ranges from 0 to 100, with higher score reflecting greater psychological resilience. CD-RISC assesses five main subdomains broadly indicating personal competence, trust/tolerance/strengthening effects of stress, acceptance of change and secure relationships, control and spiritual influences^26,29^.

Depressive behaviour was assessed through a self-administered reduced version of Patient Health Questionnaire 9^27^, translated in Italian and adapted over 8 items (hereafter called PHQ8), due to the unavailability of data on the feeling of failure. A similar measure has been reported to be robust and useful to detect depressive symptoms in population-based studies, showing operating characteristics comparable to PHQ9 in this setting^36^. This scale tests different domains related to depression, namely anhedonia, low mood, alteration of sleeping patterns or eating behaviours, feeling of failure/low self-estimate, fatigue, troubles in mental concentration, hypo-/hyperactivity behaviours, and suicidal ideation^27^. For each of these items, participants determined how much the corresponding domain had been affected in the two weeks preceding the test, which results in a score ranging from 0 (not at all) to 3 (nearly every day). Summing up these scores over the 8 items administered, the PHQ8 scale results in a total score ranging from 0 (indicating no depressive symptoms at all) to 24 (suggestive of severe depression)^36^. Of note, in this work we preferred to use PHQ8 rather than another validated reduced version of the test, over two items (PHQ2)^37^, since a larger variability of scores could provide better power to the analyses. However, the main analyses were carried out in parallel on PHQ2 to check for robustness the associations observed with PHQ8.

Mental health was tested through computing the mental component score of the validated Italian version of the self-administered SF-36 test (SF36-MCS)^38^, a widely used and thoroughly validated scale assessing health-related quality of life^28^. The questionnaire contains 36 items measuring eight multi-item parameters of health status covering the following domains: physical functioning, role limitations due to physical health problems, bodily pain, general health perceptions, vitality, social functioning, role limitations due to emotional problems and mental health. The first four domains deal with physical aspects and generate the physical component score, while the other four reflect psychological features and generate the mental component score, which was used in the present paper^28^. For each domain, a score ranking from 0 (worst health) to 100 (best health) was calculated as the weighted sum of the questions relevant to that domain. To obtain the global mental score, each mental domain was first transformed into a z-score (assigning equal weight to each item), then the resulting z-scores were combined into a global z-score through a weighted mean, using weights that resulted from a principal component analysis ^39^. Finally, the global mental health score was standardized to a normal distribution with mean = 50 and SD = 10 ^40^.

### Statistical analyses

All the scales mentioned above were analysed as continuous traits. To test the specific hypothesis that mental state or psychological factors could influence subclinical low-grade inflammation levels, which has already been proposed in the past^41–43^, we carried out stepwise generalized linear model (glm) regressions using SAS/STAT software, Version 9.4 of the SAS System for Windows© 2009. SAS Institute Inc. and SAS are registered trademarks of SAS Institute Inc., Cary, NC, USA. Four different glm models were tested for each of the psychometric scales analysed.

We first carried out a glm basic model consisting of a multivariate regression

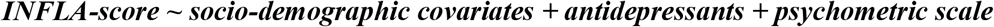

where socio-demographic covariates were age, sex and educational attainment, and antidepressants represented the reported and verified use of anti-depressants (yes/no). This model was considered as the baseline model of comparison, to partial out confounding bias of socio-demographic covariates and the effect of antidepressants on the relation between inflammation and mental health.

Then, we considered three additional multivariable regression models including potential mediators, namely disease conditions (hypertension, diabetes, cardiovascular disease, cancer, liver disease and blood disease), lifestyle factors or their proxies (smoking habits, physical activity, BMI, abdominal obesity based on waist-to-hip ratio, adherence to Mediterranean Diet and daily energy intake), and both class of mediators (full model). These factors were chosen given their reported relation with both mental health and systemic inflammation (see *Introduction* and *Discussion* sections). To allow comparability of the analyses among the different psychometric scores tested, we adjusted for the same set of covariates in all the regressions. A complete list of these covariates, including their definition, is reported in *Supplementary Methods* (Table S1a). The significance threshold for this analysis was set to α = 0.017, after applying a Bonferroni correction for three psychometric scales tested. To quantify the influence of putative mediators on the relation between INFLA-score and the psychometric scales tested, we performed a mediation analysis for each of the two classes of mediators tested in the full glm models, through the multiple mediation analysis function of the mma package in R ^44,45^, using default settings.

Then we modelled the relations between psychometric scores and each of the component biomarkers of the INFLA-score (CRP, Plt, GLR and WBC), again testing the four alternative models in SAS/STAT®. Since CRP and WBC did not attain a normal distribution, the natural logarithm of these variables was used to reduce skewness. To apply a correction for multiple testing, we performed a Principal Component Analysis (PCA) on the correlation matrices of both the three psychometric scores (Table S1b) and the four component biomarkers of INFLA-score (see Table S1c), through MatSpD (http://gump.qimr.edu.au/general/daleN/matSpD/) ^46^. This did not detect any reduction in the number of variables to correct for, hence statistical significance was set to α = 4.2×10^−3^, after applying a Bonferroni correction for four blood markers and three psychometric scores tested.

We then extended the full model glm analysis to a number of additional blood biomarkers related to inflammatory/immune response, including MPV, PCT, PDW, RDW, LY, MO, GR, NE, and NLR levels (see above for a rationale of this analysis). In this case, only the full glm model was tested, to detect residual associations with the psychometric scores analysed which were not explained by health and lifestyle mediators. For this analysis, the significance threshold was corrected for multiple testing of three different psychometric scores and eight latent blood variables, as computed on the correlation matrix of thirteen blood markers analysed (Table S1c) through MatSpD ^46^. This adjustment resulted in a final Bonferroni-corrected significance level α = 0.05 / (3×8) = 2.1×10^−3^.

## Results

Basic characteristics of the samples involved in the analyses are reported in Table 1. Compared to the samples excluded from the analysis, the datasets analysed had a higher frequency of male subjects ([49.2-51.0]% vs [45.6-46.5]%, p < 0.0001), were generally younger and more educated (p < 0.0001), showed a higher percentage of smokers (p < 0.0001) and a lower prevalence of CVD, cancer and diabetes (p < 0.0001), as well as a higher BMI (p < 0.0001). Also, they showed a higher level of physical activity (significant in PHQ8 and SF36-MCS analyses only; p < 0.001) and a higher adherence to Mediterranean diet (significant in SF36-MCS analysis only; p = 1×10^−4^). A possible explanation for these discrepancies may be that people that decide to take these self-administered psychological questionnaires are younger and more educated, since these topics are not always easily understandable and are possibly considered less important by elders and people with a lower educational level.

**Table 1.** Characteristics of the population of study, based on the psychometric score analysed. Abbreviations: CD-RISC = Connor-Davidson Resilience Scale; PHQ8 = adapted version of Patient Health Questionnaire 9 (over 8 items); SF36-MCS = 36-item Short Form Health Survey, Mental Component Score; MeDi = adherence score to Mediterranean Diet^85^. Unity of measurement is reported in brackets, where applicable. Further details on the definition of such variables are reported in *Supplementary Methods*.

| Variable | CD-RISC | PHQ8 | SF36-MCS |
| --- | --- | --- | --- |
| N | 10,415 | 12,732 | 16,952 |
| Range (min; max) | 0-100 | 0-24 | 1.81-77.91 |
| Interquartile range (Q1; Q3) | 59-75 | 1-6 | 40.67-54.33 |
| Mean, sd | 66.75 (12.42) | 3.77 (3.72) | 46.94 (9.98) |
| Median | 67 | 3 | 48.10 |
| Sex (males) | 50.96 % | 49.57 % | 49.21 % |
| Age (y; mean, sd) | 52.56 (10.71) | 53.09 (10.84) | 53.66 (11.05) |
| *Education* |  |  |  |
| Primary | 13.09 % | 14.85 % | 16.28 % |
| Lower secondary | 28.21 % | 29.70 % | 28.88 % |
| Upper secondary | 42.36 % | 40.29 % | 39.58 % |
| Post-secondary | 16.26 % | 15.06 % | 15.20 % |
| Unknown | 0.09 % | 0.09 % | 0.07 % |
| *Health conditions/*<br>*Use of specific drugs* |  |  |  |
| Hypertension | 22.89 % | 23.49 % | 24.58 % |
| Dyslipidaemia | 6.44 % | 6.61 % | 6.78 % |
| CVD | 4.27 % | 4.47 % | 4.53 % |
| Cancer | 2.73 % | 2.88 % | 2.99 % |
| Diabetes | 3.64 % | 3.68 % | 4.06 % |
| Liver Disease | 4.38 % | 4.25 % | 4.37 % |
| Blood Disease | 2.49 % | 2.46 % | 2.25 % |
| Antidepressant drugs | 2.86 % | 2.92 % | 2.83 % |
| *Lifestyle factors (or proxies)* |  |  |  |
| Smoke | 23.98 % | 24.10 % | 23.94 % |
| Energy intake<br>(Kcal/d; mean, sd) | 2147.55 (628.27) | 2140.90 (631.21) | 2132.97 (627.54) |
| MeDi score (mean, sd) | 4.34 (1.66) | 4.36 (1.66) | 4.37 (1.64) |
| Physical activity<br>(meth/d; mean, sd) | 3.43 (3.78) | 3.40 (3.78) | 3.56 (3.93) |
| BMI (kg/m<sup>2</sup>; mean, sd) | 27.54 (4.49) | 27.63 (4.54) | 27.69 (4.55) |
| Abdominal obesity | 71.29 % | 71.61 % | 71.04 % |

The results of glm regressions modelling the relation between the psychometric scales tested – namely CD-RISC, PHQ8 and SF36-MCS – and INFLA-score are reported in Table 2. Here we report standardized regression coefficients β, unless otherwise stated.

**Table 2.** Results of the stepwise glm regression of INFLA-score vs the psychometric scores analysed. Beta (β) values for standardized scales are reported, along with association p-values in brackets. Associations surviving correction for multiple testing of three psychometric scales tested (p < 0.017) are highlighted in bold. Abbreviations: COVs = covariates (confounders), including socio-demographic covariates (age, sex and educational attainment) and use of anti-depressants; MEDs = potential mediators tested; CD-RISC = Connor-Davidson Resilience Scale; PHQ8 = adapted version of Patient Health Questionnaire 9 (over 8 items); SF36-MCS = 36-item Short Form Health Survey, Mental Component Score.

| Variables | CD-RISC<br>(N=10,415) | PHQ8<br>(N=12,732) | SF36-MCS<br>(N=16,952) |
| --- | --- | --- | --- |
| Basal COVs | -0.011<br>(0.23) | **0.024**<br>**(8.9 x 10<sup>-3</sup>)** | **-0.021**<br>**(7.0 x 10<sup>-3</sup>)** |
| Basal COVs<br>+<br>Health Conditions MEDs | -0.008<br>(0.42) | **0.022**<br>**(0.015)** | -0.018<br>(0.023) |
| Basal COVs<br>+<br>Lifestyle MEDs | -0.010<br>(0.32) | -0.002<br>(0.81) | -0.014<br>(0.076) |
| Basal COVs<br>+<br>ALL MEDs | -0.007<br>(0.44) | -0.003<br>(0.78) | -0.012<br>(0.14) |

We observed significant associations of INFLA-score with PHQ8 (β = 0.024, p = 8.9×10^−3^) and SF36-MCS (β = −0.021, p = 7.0×10^−3^) in baseline models, which explained 0.48% and 0.50% of total variance in INFLA-score, respectively. These associations were attenuated after including disease conditions (β= 0.022, p = 0.015, and β = −0.018, p = 0.023), while they disappeared when lifestyle factors were added to the model (β = −0.002, p = 0.81, and β= −0.014, p = 0.076 for PHQ8 and SF36-MCS, respectively). Indeed, a multiple mediation analysis revealed that disease conditions explained 1.7% and 2.4% of the associations of INFLA-score with PHQ8 and SF36-MCS, while lifestyle factors represented 81% and 17% of the total effects, respectively. On the other hand, we observed no significant association between CD-RISC and INFLA-score, neither in the baseline model (β = −0.011, p-value = 0.23), nor after including health and lifestyle mediators in the multivariable model (see Table 2). We carried out a focused analysis of the relation between PHQ8 and SF36-MCS and the lifestyle factors (or their proxies) used in the glm models, based on the comparison of PHQ8 and SF36-MCS distribution quartiles in the datasets analysed (Tables S2a-c and S3a-c). We observed that a higher PHQ8 was associated with a higher frequency of smoking (χ^2^ = 51.3, p < 10^−4^), BMI (one-way ANOVA F = 13.8, p < 10^−4^) and prevalence of abdominal obesity (χ^2^ = 17.6, p = 5×10^−4^), as well as with lower adherence to Mediterranean Diet (F = 5.7, p = 7×10^−4^), energy intake (F = 9.8, p < 10^−4^) and energy expenditure in leisure physical activity (F = 50.0, p < 10^−4^). In line with these observations, subjects with better SF36-MCS scores showed a lower frequency of smokers (χ^2^ = 68.9, p < 10^−4^), and higher adherence to Mediterranean Diet (F = 13.1, p < 10^−4^), daily energy intake (F = 13.7, p < 10^−4^) and physical activity (F = 78.9, p = 3.1×10^−4^). Upper SF36-MCS quartiles also showed a higher BMI (F = 4.6, p = 9×10^−4^) and a non-significant higher frequency of abdominal obesity (χ ^2^= 4.8, p = 0.19).

When we modelled the relation of the psychometric scores analysed with each of the four components of INFLA-score, we observed no significant associations in the baseline models surviving correction for multiple testing (p > 4.2×10^−3^), with the exception of the models involving GLR (Tables S4a-d). Indeed, for this biomarker we observed significant baseline associations with all of the three psychometric scores tested (positive with PHQ8, negative with CD-RISC and SF36-MCS; see Table S4d). For one of these scores, SF36-MCS, the association remained statistically significant even after including all the mediators tested in the full model (β = −0.033, p < 10^−4^), while it was only nominally significant for PHQ8 (β = 0.018, p = 0.01). In all cases, associations were generally attenuated by lifestyle mediators (Table S4d). Full glm models on a number of circulating biomarkers of low-grade inflammation revealed additional significant associations surviving correction for multiple testing (p < 2.1×10^−3^; see Table 3). The most significant associations were observed for PDW with PHQ8 (β = 0.059, p = 2×10^−4^) and CD-RISC (β = −0.53, p = 2×10^−3^), as well as between SF36-MCS and MO (β = 0.065, p < 10^−4^), LY (β = 0.3, p < 10^−4^), GR (β = −0.37, p < 10^−4^), NE(β = −0.35, p < 10^−4^), GLR (β = −0.032, p < 10^−4^) and NLR (β = −0.034, p < 10^−4^).

**Table 3.**
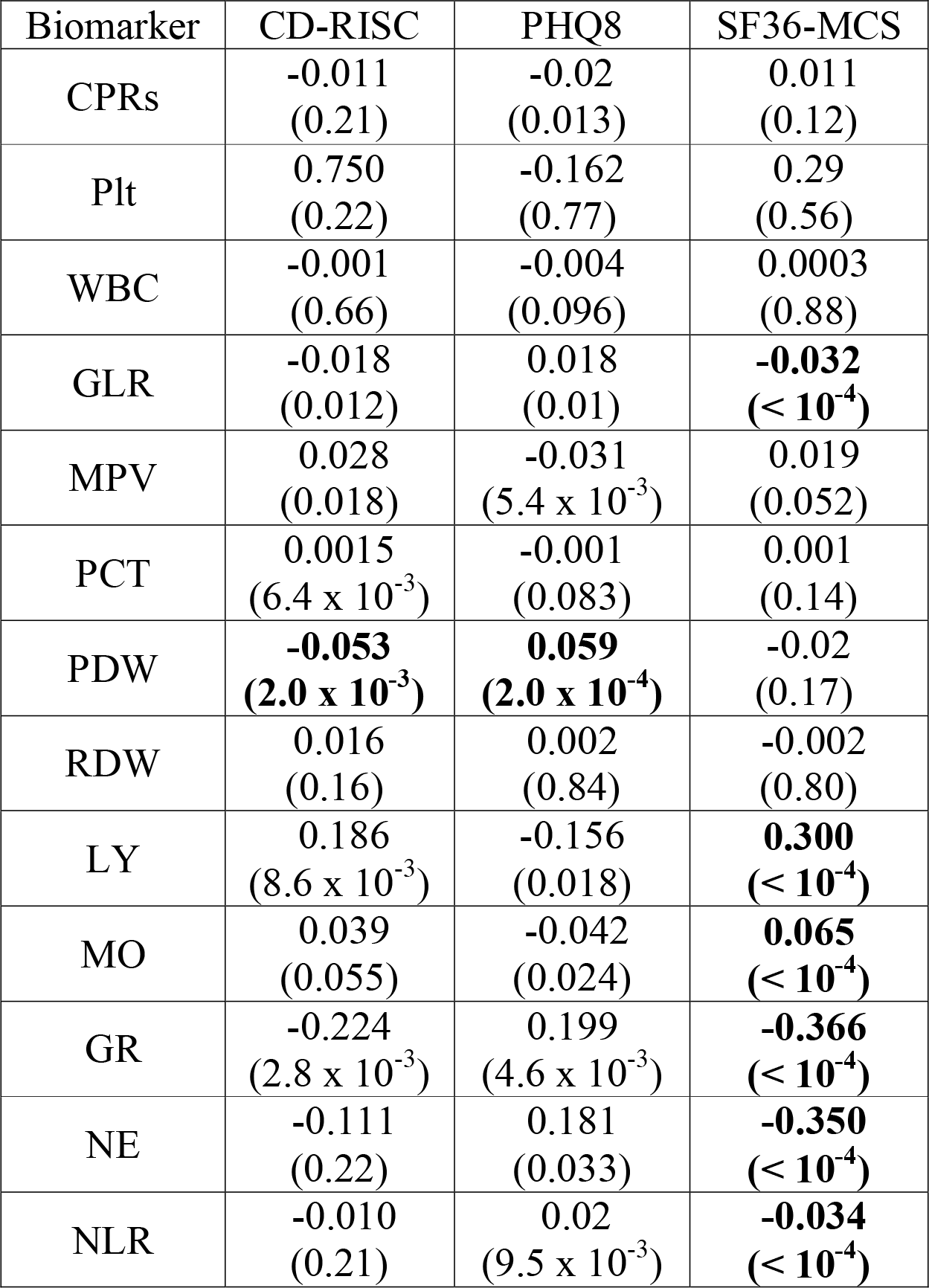
Results of the “full” glm analysis investigating the relation between the psychometric scores tested and a number of inflammation-related circulating biomarkers. Associations surviving correction for multiple testing (p < 2.1 × 10^−3^) are highlighted in bold. Abbreviations: CRP = C-reactive protein (mg/L, log scale); Plt = platelet count (×10^9^/L); WBC = white blood cell count (×10^9^/L, log scale); GLR and NLR = granulocyte- and neutrophil-to-lymphocyte ratio; MPV = mean platelet volume (fL); PCT = plateletcrit (%); PDW = platelet distribution width (%); RDW = red cell distribution width (%); LY, MO, GR and NE = fraction of lymphocytes, monocytes, granulocytes and neutrophils over the total WBC (%).

| Biomarker | CD-RISC | PHQ8 | SF36-MCS |
| --- | --- | --- | --- |
| CPRs | -0.011<br>(0.21) | -0.02<br>(0.013) | 0.011<br>(0.12) |
| Plt | 0.750<br>(0.22) | -0.162<br>(0.77) | 0.29<br>(0.56) |
| WBC | -0.001<br>(0.66) | -0.004<br>(0.096) | 0.0003<br>(0.88) |
| GLR | -0.018<br>(0.012) | 0.018<br>(0.01) | **-0.032**<br>**(< 10<sup>-4</sup>)** |
| MPV | 0.028<br>(0.018) | -0.031<br>(5.4 x 10<sup>-3</sup>) | 0.019<br>(0.052) |
| PCT | 0.0015<br>(6.4 x 10<sup>-3</sup>) | -0.001<br>(0.083) | 0.001<br>(0.14) |
| PDW | **-0.053**<br>**(2.0 x 10<sup>-3</sup>)** | **0.059**<br>**(2.0 x 10<sup>-4</sup>)** | -0.02<br>(0.17) |
| RDW | 0.016<br>(0.16) | 0.002<br>(0.84) | -0.002<br>(0.80) |
| LY | 0.186<br>(8.6 x 10<sup>-3</sup>) | -0.156<br>(0.018) | **0.300**<br>**(< 10<sup>-4</sup>)** |
| MO | 0.039<br>(0.055) | -0.042<br>(0.024) | **0.065**<br>**(< 10<sup>-4</sup>)** |
| GR | -0.224<br>(2.8 x 10<sup>-3</sup>) | 0.199<br>(4.6 x 10<sup>-3</sup>) | **-0.366**<br>**(< 10<sup>-4</sup>)** |
| NE | -0.111<br>(0.22) | 0.181<br>(0.033) | **-0.350**<br>**(< 10<sup>-4</sup>)** |
| NLR | -0.010<br>(0.21) | 0.02<br>(9.5 x 10<sup>-3</sup>) | **-0.034**<br>**(< 10<sup>-4</sup>)** |

Associations between PHQ2, INFLA-score and its component biomarkers were substantially consistent with those observed with PHQ8 (Table S5a), as were the associations with the different inflammation-related biomarkers tested above (Table S5b).

## Discussion

In this paper, we investigated the relation between subclinical low-grade inflammation and three psychometric scores in the Italian population, representing distinct but correlated psychological domains linked to mental health: psychological resilience, depressive symptoms and mental wellbeing. We did this through a step-wise approach.

First we tested associations between mental health and low-grade inflammation, which revealed a significant positive association of depressive symptoms (PHQ8) and a negative association of mental wellbeing (SF36-MCS) with INFLA-score. These findings are in line with previous studies suggesting an association of inflammation with depression (reviewed in ^6,7^) and with measures of quality of life, both in physical and in mental domains^13–18^. The above mentioned associations were attenuated by including lifestyle factors as covariates in the model, which explained 81% and 17% of the associations with PHQ8 and SF36-MCS, respectively. This evidence suggests an important role of lifestyle in mediating the effect of depressive symptoms and mental wellbeing on low-grade inflammation. Likely, subjects presenting with depressive symptoms or impaired mental wellbeing worsen their lifestyle, e.g. avoiding to practice physical activity on a regular basis or decreasing their adherence to healthy dietary patterns. Our data corroborate this hypothesis, showing that subjects with better mental health – i.e. those in the lower quartiles of PHQ8 and in the upper quartiles of SF36-MCS distribution – report a higher adherence to Mediterranean diet, a higher level of physical activity, and a lower tendency to smoke.

Physical exercise has been investigated in relation to mental health disorders, with studies consistently reporting a beneficial effect on depressive symptoms, stress, anxiety and suicidality^47–49^. Several studies, including clinical trials, support the idea that physical activity may have both preventive and treatment effects on depression and anxiety disorders^47,50^. On the other hand, physical activity has been associated with lower levels of inflammatory markers, in an independent way from other factors favouring inflammation, like obesity^51^.

Diet has recently gained attention as a potential target for the treatment of psychiatric disorders, giving rise to the new field of “Nutritional Psychiatry”^52^. Meta-analyses have reported an inverse relation between healthy dietary patterns, especially Mediterranean Diet and reduced depression risk^53–55^. This association seems to be independent of other key health behaviours discussed here, including physical activity and smoking, and of environmental factors like socioeconomic status. Moreover, it is consistent across countries, cultures and age groups^52,54^. Of note, our group previously reported positive associations of adherence to Mediterranean Diet with psychological resilience^29^ and mental wellbeing^56^ in the Moli-sani study. Interestingly, a large longitudinal study detected an increased depression risk among women following a pro-inflammatory dietary pattern^57^, an evidence supported by a recent meta-analysis^54^, and by the finding that adherence to Mediterranean Diet is associated with the methylation of genes reportedly related to inflammation^58^.

Epidemiological studies on smoking habits and depression consistently suggest that depressive symptoms and mental disorders are associated with higher levels of cigarette consumption, and people with depression appear to have more difficulties in quitting smoking^59^. In this view, depression and low mood appear to be a barrier to smoking cessation and sustained abstinence, and are associated with increased risk of smoking relapse in the long-term. On the other hand, quitting smoking may have mental health benefits and is considered to lower depression risk^59^. Similarly, even mental wellbeing as measured in our paper has been associated with lower odds of smoking initiation and higher odds of successful smoking cessation^60^. Again, smoking has been previously associated with the circulating levels of inflammatory markers, both in cross-sectional^61,62^ and in longitudinal studies^63^. Overall, these findings suggest a potential role of the main lifestyle factors in modulating the relationship between mental health and inflammation, in line with the evidence reported in this paper.

Among the four component markers of the INFLA-score, only GLR showed evidence of association with the psychometric scores analysed, which was only slightly attenuated by the adjustment for health conditions and lifestyle mediators. This suggests that the specific relation between mental wellbeing and GLR could also be explained by other, likely biological, factors (see below for further discussion). This evidence extended to resilience and depressive symptoms, and to other circulating biomarkers which are involved in inflammatory processes but were not included in the INFLA-score: PDW, LY, MO, GR, NE and NLR. Interestingly, these findings are in line with a recent blood transcriptomic analysis which identified a set of 165 differentially expressed genes between MDD cases and healthy controls, in two independent datasets^64^. Of these, 90 genes which were over-expressed in MDD were significantly enriched for Gene Ontology terms related to innate immunity and included clusters of genes with correlated expression in monocytes, monocyte-derived dendritic cells, and neutrophils. On the other hand, 75 genes under-expressed in MDD were associated with adaptive immunity and included clusters of genes with correlated expression in lymphocytes and erythroblasts^64^. Additional findings are in line with our results, such as the activation of the innate immune system and increased pro-inflammatory cytokine signalling in depressed subjects^65,66^, and the co-occurrence of lymphocyte deficiencies and inflammatory monocyte activation as related phenomena in the same patients with MDD^67–70^.

Among the putative biological factors which may explain the residual associations observed between inflammation markers and mental health, a prominent role may have genetic factors like Single Nucleotide Polymorphisms (SNPs), Copy Number Variants (CNVs), or rare point mutations. However, previous Genome-Wide Association Studies (GWAS) have reported a lack of genetic correlations between MDD risk, Plt and MPV^71^, and non-significant associations in multivariable Mendelian Randomization (MR) analyses carried out between MDD risk and a number of blood cell indexes which overlap or are largely correlated with those tested here^72^. Although these results warrant a deeper analysis, they pave the way to the hypothesis that alternative biological factors, e.g. epigenetic modifications, may regulate the expression of inflammation-related genes^73^. This hypothesis is particularly interesting in light of the above-mentioned transcriptomic findings by Leday et al.^64^, and of the increasing role attributed to epigenetic changes in the pathophysiological mechanism of mental health and depression, in response to environmental stressors (reviewed in ^74,75^). Environmental factors may also mediate the relation between mental health and low-grade inflammation, either influencing epigenetic modifications or through alternative/complementary mechanisms, like gene-by-environment (GxE) interactions. Similar mechanisms are suggested to have a prominent role in the aetiology of depression and of other psychiatric disorders^76^, and may also influence the link with low-grade inflammation.

In this study, we modelled the relation between subclinical inflammation and mental status hypothesizing that the latter could influence the former. Although there is contrasting evidence on the directionality of this relationship^41–43,77–79^, one of these studies supported a bi-directional effect between mental health and inflammation^79^. To our view, this represents the most likely scenario, where both psychological and inflammatory status may reciprocally amplify their effects through a positive feedback mechanism (Figure 1). Consistent with this, it has been hypothesized that cytokines may promote depression, which in turn would upregulate inflammatory signalling^4^. In this view, it would be important to break this pathological loop through effective interventions aimed at lowering the systemic inflammation levels, which could be achieved by acting on the lifestyle factors discussed above.

**Figure 1.**
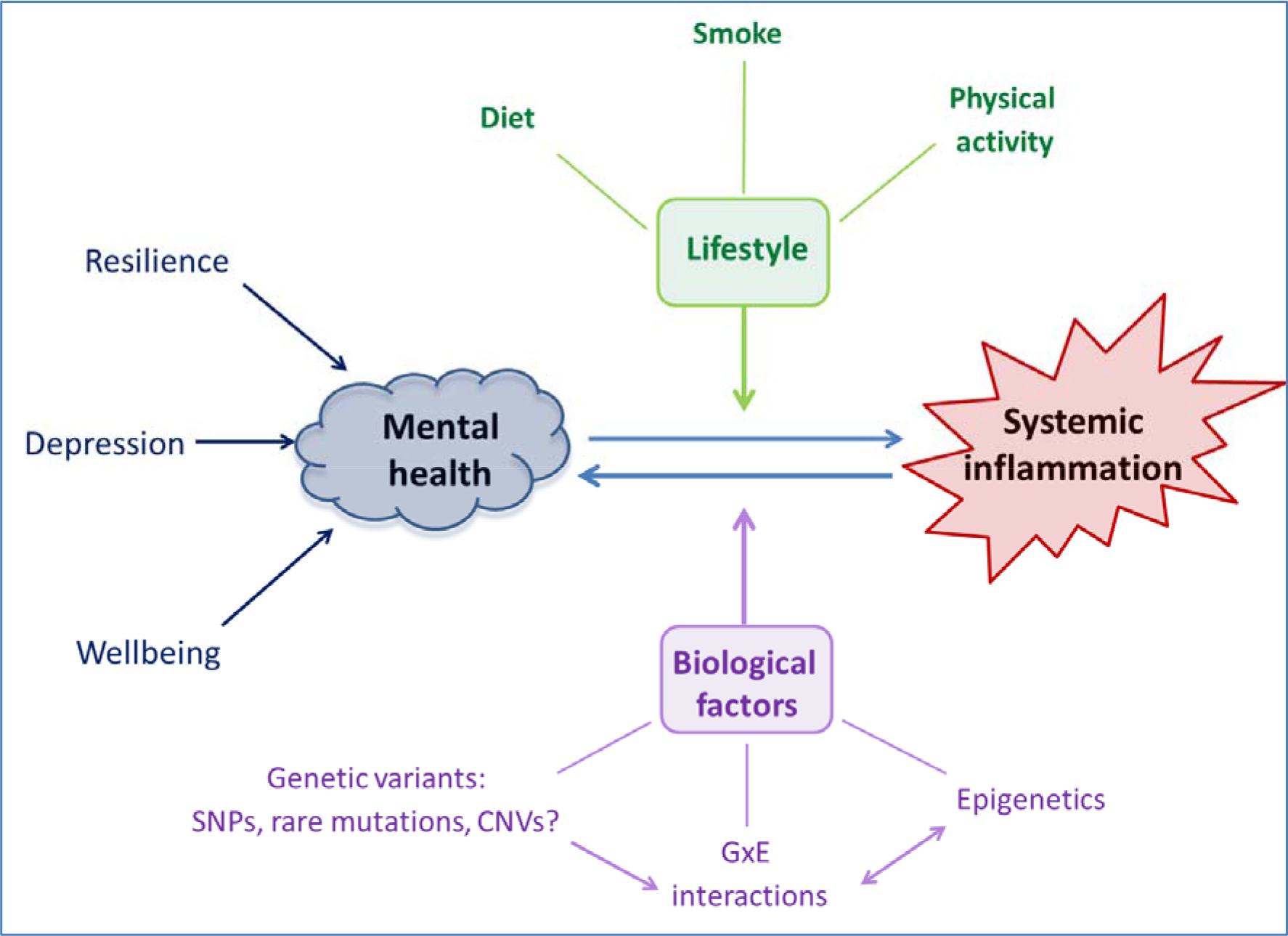
Hypothesized bi-directional relation between mental health and systemic inflammation. Main environmental and biological factors potentially mediating this association are also reported. Abbreviations: SNPs = Single Nucleotide Polymorphisms; CNVs = Copy Number Variants; GxE = Gene-by-Environment interactions.

### Strengths and limitations

Our study presents different points of strength, including the availability of a rich set of blood markers in an unselected general population cohort, with a relatively large sample size (N_max_~17,000), and a well characterized dataset in terms of socio-economic, lifestyle, anthropometric and medical information.

The main limitation of the present study is represented by the current unavailability of longitudinal data, which does not allow us to make causal inference on the relation between inflammation and mental health. However, a recall of the cohort is currently ongoing, in which all the psychometric scales tested in the present paper are being collected. This active follow-up will allow us to tackle another potential limitation of this study, namely the use of a non-validated scale to assess depressive symptoms (PHQ8, see *Methods* section), which is now replaced by a validated test (PHQ9). However, parallel analyses on a reduced validated scale for screening of depressive symptoms (PHQ2) revealed results consistent with the main analysis on PHQ8.

Similarly, we used here a non-validated INFLA-score, with the aim of tapping into the different domains of systemic inflammation. Since low-grade inflammation is a condition not yet consistently defined or measured^80^, and no established definition or cut-off exist, the lack of a gold standard measure makes it challenging to validate a score aimed at assessing a sub-clinical inflammation status. However, independent lines of evidence support the fact that the INFLA-score actually provides an estimation of low-grade inflammation. First, INFLA-score resulted associated with known conditions/behaviours related to inflammation. We recently demonstrated the association between INFLA-score with a dietary inflammation index^81^, and previously showed that the INFLA-score is directly associated with pro-inflammatory health conditions (i.e. obesity, abdominal adiposity), or behaviours (smoking, low physical exercise)^82^. We replicated these associations in an independent family-based cohort from the Molise region (N=745; Bonaccio, personal communication). Moreover, the INFLA-score is positively associated with the levels of soluble P-selectin, which is involved in the inflammatory process^83^. Last, the INFLA-score was also found negatively associated with methylation levels in the *PEAR1* gene, in two independent cohorts from Italy and Belgium (Izzi et al., under review).

Finally, our study leaves an open question on which biological factors influence the residual association between inflammation-related markers and the psychometric scales analysed. Although we might be able to answer this question only when genetic data will be available in our cohort, here we tried to speculate on this hypothesis based on the results of genetic analyses previously published.

## Conclusion

Overall, this study underlines the importance of lifestyle factors as a whole in the relation between mental health and low-grade inflammation, supporting them as a therapeutic target not only for the treatment of mental disorders^84^, but also to lower the risk of comorbid conditions such as CVD, cancer and diabetes. These factors represent therapeutic goals easier to target than the “biological” ones, through treatments which imply fewer side effects and a lower cost-to-benefit ratio. On the other hand, our data suggest that biological factors are implicated in this relation, underlining the potential of genetic and epigenetic research on mental health and disorders to enlighten their link with chronic health conditions.

## Acknowledgements and Disclosures

The Moli-sani Research Investigators thank the Associazione Cuore Sano Onlus (Campobasso, Italy) for its cultural and financial support. The enrolment phase of the Moli-sani Study was supported by research grants from Pfizer Foundation (Rome, Italy), the Italian Ministry of University and Research (MIUR, Rome, Italy) – Programma Triennale di Ricerca, Decreto no. 1588 and Instrumentation Laboratory, Milan, Italy. The present analyses were partially supported by the Italian Ministry of Health 2013 (Young investigator grant to MB, no. GR-2013-02356060) and by the Italian Association for Cancer Research (AIRC) with grant AIRC ‘5 ×1000’ to LI, Ref. no. 12237. Funders had no role in this study design, collection, analysis, and interpretation of data, nor in the writing and submission phase of the manuscript. MB was supported by a Fondazione Umberto Veronesi Fellowship. SC was the recipient of a Fondazione Umberto Veronesi Travel Grant. AG, MB, LI and GdG contributed to the conception and design of the study, supported by CC and MBD. SC and ADC managed data collection of the Moli-sani Study. MS designed and was responsible for the psychometric assessment in the study. AG, MB, ADC and SC analysed the data; AG wrote the first draft of paper, with contributions and critical review from all the co-authors. LI, MBD, ADC, CC and GdG originally inspired the Moli-sani Study. We thank Dr Benedetta Izzi for a critical read of the paper. All Authors were and are independent from funders and declare no conflicts of interest.

## Moli-sani Study Investigators

The enrolment phase of the Moli-sani Study was conducted at the Research Laboratories of the Catholic University in Campobasso (Italy), the follow up of the Moli-sani cohort is being conducted at the Department of Epidemiology and Prevention of the IRCCS Neuromed, Pozzilli, Italy. *Steering Committee:* Licia Iacoviello (Chairperson), Giovanni de Gaetano and Maria Benedetta Donati. *Scientific secretariat:* Licia Iacoviello (Coordinator), Marialaura Bonaccio, Americo Bonanni, Chiara Cerletti, Simona Costanzo, Amalia De Curtis, Giovanni de Gaetano, Augusto Di Castelnuovo, Maria Benedetta Donati, Francesco Gianfagna, Mariarosaria Persichillo, Teresa Di Prospero (Secretary). *Safety and Ethycal Committee:* Jos Vermylen (Catholic Univesity, Leuven, Belgium) (Chairperson), Ignacio De Paula Carrasco (Accademia Pontificia Pro Vita, Roma, Italy), Simona Giampaoli (Istituto Superiore di Sanità, Roma, Italy), Antonio Spagnuolo (Catholic University, Roma, Italy). *External Event adjudicating Committee:* Deodato Assanelli (Brescia, Italy), Vincenzo Centritto (Campobasso, Italy). *Baseline and Follow-up data management:* Simona Costanzo (Coordinator), Marco Olivieri (Università del Molise, Campobasso, Italy). *Informatics:* Marco Olivieri (Università del Molise, Campobasso, Italy). *Data Analysis:* Augusto Di Castelnuovo (Coordinator), Marialaura Bonaccio, Simona Costanzo, Alessandro Gialluisi, Francesco Gianfagna, Emilia Ruggiero. *Biobank and biomedical analyses:* Amalia De Curtis (Coordinator), Sara Magnacca. *Genetic analyses:* Benedetta Izzi (Coordinator), Francesco Gianfagna, Claudio Grippi, Annalisa Marotta, Fabrizia Noro. *Communication and Press Office:* Americo Bonanni (Coordinator), Francesca De Lucia (Associazione Cuore Sano, Campobasso, Italy). *Recruitment staff:* Mariarosaria Persichillo (Coordinator), Francesca Bracone, Francesca De Lucia (Associazione Cuore Sano, Campobasso, Italy), Salvatore Dudiez, Livia Rago. *Follow-up Event adjudication:* Livia Rago (Coordinator), Simona Costanzo, Amalia De Curtis, Licia Iacoviello, Teresa Panzera, Mariarosaria Persichillo. *Regional Health Institutions:* Direzione Generale per la Salute - Regione Molise; Azienda Sanitaria Regionale del Molise (ASReM, Italy); Molise Dati Spa (Campobasso, Italy); Offices of vital statistics of the Molise region. *Hospitals:* Presidi Ospedalieri ASReM: Ospedale A. Cardarelli – Campobasso, Ospedale F. Veneziale – Isernia, Ospedale San Timoteo - Termoli (CB), Ospedale Ss. Rosario - Venafro (IS), Ospedale Vietri – Larino (CB), Ospedale San Francesco Caracciolo - Agnone (IS); Casa di Cura Villa Maria - Campobasso; Fondazione di Ricerca e Cura Giovanni Paolo II - Campobasso; IRCCS Neuromed - Pozzilli (IS). Baseline Recruitment staff is available at http://www.moli-sani.org/index.php?option=com_content&task=view&id=21128&Itemid=118

