## Supplemental Data for "Lifestyle and biological factors influence the relationship between mental health and low-grade inflammation"

**Supplementary Methods**

***Blood measurements***

Blood samples were obtained between 7 and 9 AM from participants who had fasted overnight and had refrained from smoking for at least 6 h. Biochemical analyses were performed in the centralised Moli-sani laboratory ^1^. All haemochromocytometric analyses (white blood cell and platelet counts, granulocyte % and lymphocyte %) were performed by cell counter (Coulter HMX, Beckman Coulter, Milan, Italy) within 3 hours (h) from venipuncture. Since the measurements of neutrophil % showed a high missing rate, granulocyte % was considered as an accurate predictor and hence the granulocyte to lymphocyte ratio (GLR) was preferred for the computation of the INFLA-score ^2^. High sensitivity C reactive protein (CRP) was measured in fresh serum, by a latex particle-enhanced immunoturbidimetric assay (ILab 350 Instrumentation Laboratory, Milan, Italy).

***Definition of covariates and potential mediators***

All the variables explained below were used in the analysis as continuous variables, unless otherwise stated.

For e*ducational attainment*, subjects were divided into four categories, based on their education level (i.e. the school level that they had completed): *Primary, Lower secondary, Upper secondary* and *Post-secondary*.

The covariate *antidepressants* was based on the self-reported use of antidepressant drugs and coded as a binary variable (Yes/No). This was further verified by the recruitment staff by asking the participants to show the drug box in use, and only those subjects which were able to provide such a proof were classified as antidepressant users.

Prevalent *hypertension, dyslipidaemia* and *diabetes* were defined as dichotomous variables (Yes/No), based on the reported and verified use of specific drugs for the treatment of these disorders.

Prevalent *cardiovascular disease* (*CVD*) and *cancer* were defined as three-classes variables: subjects with no medical history of the disease, subjects reporting the disease with medical documentation and those reporting the disease without medical documentation.

*Liver disease* and *blood disease* were defined as dichotomous variables, based on the report of being affected by the disease or not.

For *smoking status*, subjects were assigned to three categories based on their cigarette smoking habits: *smokers, ex-smokers* (i.e. subjects who quitted at least one year before the interview) and *non-smokers*.

Leisure-time *physical activity* was assessed through a structured questionnaire and expressed as daily energy expenditure in metabolic equivalent task-hours (MET-h/day) ^3^.

Food intake was assessed by the validated Italian EPIC food frequency questionnaire ^4^. Adherence to *Mediterranean diet* was defined according to the Mediterranean Diet Score ^5^. The EPIC questionnaire also allowed to compute the daily *energy intake* for the subjects assessed (Kcal/day).

Height and weight were measured for each participant and *body mass index (BMI)* was calculated as kg/m^2^. Waist circumference (cm) was measured in the middle between the 12^th^ rib and the iliac crest, while hip circumference (cm) was measured around the buttocks. Then waist-to-hip ratio (WHR) was calculated, and the resulting measure of *abdominal obesity* was inferred as a dichotomous variable (Yes/No), based on different thresholds for the two sexes: subjects were defined as obese if they had WHR ≥ 0.90 and ≥ 0.85 for men and women, respectively ^6^. Both BMI and abdominal obesity, which are measures of relative weight and fat distribution, were used here as proxies of dietary patterns and physical activity.

| Covariates and Mediators | CD-RISC | PHQ8 | SF36-MCS |
| --- | --- | --- | --- |
| Basal covariates | age + sex + educational attainment + use of antidepressants | | |
| Health Conditions  Mediators | hypertension + dyslipidaemia + cancer + CVD + diabetes + liver disease + blood disease | | |
| Lifestyle  Mediators | smoke + leisure + BMI + abdominal obesity + MeDi + calory intake | | |

**Table S1a.** Covariates and mediators used in the generalised linear model (glm) regressions. Legend: CD-RISC = Connor-Davidson Resilience Scale; PHQ8 = adapted version of Patient Health Questionnaire 9 (over 8 items); SF36-MCS = 36-item Short Form Health Survey, Mental Component Score.

**b)**

| Scale | CD-RISC | PHQ8 | SF36-MCS |
| --- | --- | --- | --- |
| CD-RISC | 1 | -0.244 | 0.314 |
| PHQ8 | <.0001 | 1 | -0.645 |
| SF36-MCS | <.0001 | <.0001 | 1 |

**c)**

| Biomarker | CRP | Plt | MPV | PCT | PDW | RDW | WBC | LY | MO | GR | NE | GLR | NLR |
| --- | --- | --- | --- | --- | --- | --- | --- | --- | --- | --- | --- | --- | --- |
| CRP | 1 | 0.075 | -0.006 | 0.072 | -0.002 | 0.049 | 0.247 | -0.106 | -0.024 | 0.107 | 0.103 | 0.11 | 0.11 |
| Plt | <.0001 | 1 | -0.267 | 0.85 | -0.151 | 0.1 | 0.231 | -0.042 | -0.021 | 0.045 | 0.06 | 0.044 | 0.051 |
| MPV | 0.35 | <.0001 | 1 | 0.257 | -0.367 | 0.261 | 0.022 | 0.034 | 0.116 | -0.064 | -0.062 | -0.045 | -0.064 |
| PCT | <.0001 | <.0001 | <.0001 | 1 | -0.369 | 0.245 | 0.245 | -0.022 | 0.037 | 0.011 | 0.023 | 0.018 | 0.012 |
| PDW | 0.74 | <.0001 | <.0001 | <.0001 | 1 | -0.197 | -0.02 | -0.078 | -0.098 | 0.1 | 0.097 | 0.086 | 0.105 |
| RDW | <.0001 | <.0001 | <.0001 | <.0001 | <.0001 | 1 | 0.018 | -0.048 | 0.004 | 0.044 | 0.029 | 0.05 | 0.037 |
| WBC | <.0001 | <.0001 | 0.001 | <.0001 | 0.003 | 0.01 | 1 | -0.237 | -0.178 | 0.271 | 0.27 | 0.258 | 0.26 |
| LY | <.0001 | <.0001 | <.0001 | 0.001 | <.0001 | <.0001 | <.0001 | 1 | 0.073 | -0.963 | -0.938 | -0.991 | -0.987 |
| MO | 0.0004 | 0.002 | <.0001 | <.0001 | <.0001 | 0.56 | <.0001 | <.0001 | 1 | -0.341 | -0.341 | -0.174 | -0.194 |
| GR | <.0001 | <.0001 | <.0001 | 0.12 | <.0001 | <.0001 | <.0001 | <.0001 | <.0001 | 1 | 0.972 | 0.982 | 0.982 |
| NE | <.0001 | <.0001 | <.0001 | 0.004 | <.0001 | 0.0004 | <.0001 | <.0001 | <.0001 | <.0001 | 1 | 0.956 | 0.975 |
| GLR | <.0001 | <.0001 | <.0001 | 0.008 | <.0001 | <.0001 | <.0001 | <.0001 | <.0001 | <.0001 | <.0001 | 1 | 0.995 |
| NLR | <.0001 | <.0001 | <.0001 | 0.13 | <.0001 | <.0001 | <.0001 | <.0001 | <.0001 | <.0001 | <.0001 | <.0001 | 1 |

**Table S1.** Correlation matrix of **b)** three psychometric scales and **c)** thirteen blood markers tested. Pearson´s r correlation coefficients are reported in the upper triangle, with relevant p-values in the lower triangle. Abbreviations: CD-RISC = Connor-Davidson Resilience Scale; PHQ8 = adapted version of Patient Health Questionnaire 9 (over 8 items); SF36-MCS = 36-item Short Form Health Survey, Mental Component Score.

**Supplementary Results**

**a)**

| **Quartile** | **PHQ8**  **score** | **N** | **BMI**  **(kg/m^2^)** | **MeDi**  **Trichopulou score** | **Energy Intake**  **(Kcal/day)** | **Physical Activity**  **(MET-h/day)** |
| --- | --- | --- | --- | --- | --- | --- |
| **Q1** | 0 | 2663 | 0.91  (0.07) | 4.4  (1.64) | 2184.64  (631.03) | 3.9  (3.86) |
| **Q2** | 1-2 | 3253 | 0.91  (0.08) | 4.4  (1.68) | 2156.48  (626.31) | 3.68  (3.98) |
| **Q3** | 3-5 | 3431 | 0.91  (0.08) | 4.38  (1.65) | 2131.78  (614.85) | 3.33  (3.79) |
| **Q4** | 6-24 | 3385 | 0.91  (0.08) | 4.26  (1.66) | 2100.76  (649.7) | 2.81  (3.4) |

**b)**

| **Smoking status** | **Q1** | **Q2** | **Q3** | **Q4** |
| --- | --- | --- | --- | --- |
| **non-smokers** | 1281 | 1452 | 1709 | 1668 |
| **ex-smokers** | 785 | 1006 | 936 | 827 |
| **current smokers** | 597 | 795 | 786 | 890 |

**c)**

| **Abdominal Obesity** | **Q1** | **Q2** | **Q3** | **Q4** |
| --- | --- | --- | --- | --- |
| **no** | 817 | 966 | 930 | 901 |
| **yes** | 1846 | 2287 | 2501 | 2484 |

**Table S2.** Descriptives of lifestyle variables according to quartiles of PHQ8 distribution. We report **a)** mean (SD) values of BMI, adherence score to Mediterranean Diet (MeDi)^5^, daily energy intake (Kcal/day) and expenditure in physical activity (MET-h/day), as well as contingency tables of **b)** smoking status and **c)** abdominal obesity by PHQ8 quartiles.

**a)**

| **Quartile** | **SF36-MCS score** | **N** | **BMI**  **(kg/m^2^)** | **MeDi**  **Trichopulou score** | **Energy Intake**  **(Kcal/day)** | **Physical Activity**  **(MET-h/day)** |
| --- | --- | --- | --- | --- | --- | --- |
| **Q1** | 1.81-40.67 | 4238 | 27.52  (4.63) | 4.26  (1.66) | 2084.52  (630.42) | 2.86  (3.4) |
| **Q2** | 40.68-48.10 | 4238 | 27.72  (4.62) | 4.34  (1.62) | 2130.48  (615.07) | 3.46  (3.98) |
| **Q3** | 48.11-54.33 | 4238 | 27.63  (4.5) | 4.43  (1.66) | 2149.29  (624.64) | 3.81  (3.99) |
| **Q4** | 54.34-77.91 | 4238 | 27.87  (4.43) | 4.46  (1.63) | 2167.58  (636.99) | 4.1  (4.2) |

**b)**

| **Smoking status** | **Q1** | **Q2** | **Q3** | **Q4** |
| --- | --- | --- | --- | --- |
| **non-smokers** | 2173 | 2010 | 2003 | 1948 |
| **ex-smokers** | 999 | 1186 | 1254 | 1321 |
| **current smokers** | 1066 | 1042 | 981 | 969 |

**c)**

| **Abdominal Obesity** | **Q1** | **Q2** | **Q3** | **Q4** |
| --- | --- | --- | --- | --- |
| **no** | 1259 | 1260 | 1201 | 1190 |
| **yes** | 2979 | 2978 | 3037 | 3048 |

**Table S3.** Descriptives of lifestyle variables according to quartiles of SF36-MCS distribution. We report **a)** mean (SD) values of BMI, adherence score to Mediterranean Diet (MeDi)^5^, daily energy intake (Kcal/day) and expenditure in physical activity (MET-h/day), as well as contingency tables of **b)** smoking status and **c)** abdominal obesity by SF36-MCS quartiles.

**a)**

| Variables | CD-RISC | PHQ8 | SF36-MCS |
| --- | --- | --- | --- |
| Basal COVs | -0.002  (0.82) | 0.009  (0.29) | 0.012  (0.11) |
| Basal COVs  +  Health Conditions MEDs | -0.002  (0.87) | 0.006  (0.51) | 0.015  (0.038) |
| Basal COVs  +  Lifestyle MEDs | -0. 010  (0.22) | -0.019  (0.017) | 0.010  (0.13) |
| Basal COVs  +  ALL MEDs | -0.011  (0.21) | -0.020  (0.013) | 0.011  (0.12) |

**b)**

| Variables | CD-RISC | PHQ8 | SF36-MCS |
| --- | --- | --- | --- |
| Basal COVs | 0.67  (0.27) | -0.208  (0.71) | 0.403  (0.41) |
| Basal COVs  +  Health Conditions MEDs | 0.72  (0.24) | 0.013  (0.98) | 0.264  (0.59) |
| Basal COVs  +  Lifestyle MEDs | 0.69  (0.26) | -0.405  (0.47) | 0.442  (0.37) |
| Basal COVs  +  ALL MEDs | 0.75  (0.22) | -0.162  (0.77) | 0.285  (0.56) |

**c)**

| Variables | CD-RISC | PHQ8 | SF36-MCS |
| --- | --- | --- | --- |
| Basal COVs | -0.003  (0.15) | 0.004  (0.088) | -0.004  (0.044) |
| Basal COVs  +  Health Conditions MEDs | -0.002  (0.30) | 0.003  (0.15) | -0.003  (0.13) |
| Basal COVs  +  Lifestyle MEDs | -0.002  (0.39) | -0.003  (0.15) | -0.0007  (0.71) |
| Basal COVs  +  ALL MEDs | -0.001  (0.66) | -0.004  (0.096) | 0.0003  (0.89) |

**d)**

| Variables | CD-RISC | PHQ8 | SF36-MCS |
| --- | --- | --- | --- |
| Basal COVs | -0.023  (**1.4 x 10^-3^**) | **0.022**  **(1.0 x 10^-3^)** | **-0.042**  **(< 10^-4^)** |
| Basal COVs  +  Health Conditions MEDs | -0.023  (**1.4 x 10^-3^**) | **0.020**  **(3.1 x 10^-3^)** | **-0.040**  **(< 10^-4^)** |
| Basal COVs  +  Lifestyle MEDs | -0.019  (0.011) | **0.020**  **(3.2 x 10^-3^)** | **-0.036**  **(< 10^-4^)** |
| Basal COVs  +  ALL MEDs | -0.018  (0.012) | 0.018  (0.01) | **-0.033**  **(< 10^-4^)** |

**Table S4.** Results of the stepwise regressions for the four component markers of the INFLA-score, **a)** C-reactive protein (CRP; mg/L, log scale), **b)** platelets count (Plt; x10^9^/L), **c)** white blood cells count (WBC; x10^9^/L, log scale) and **d)** granulocyte-to-lymphocite ratio (GLR) vs the psychometric scores analysed. ^a^ Beta (β) values referring to standardized psychometric scales (but not standardized markers) are reported, along with association p-values in brackets. Abbreviations: COVs = covariates (confounders, see Table S1a above); MEDs = potential mediators tested; CD-RISC = Connor-Davidson Resilience Scale; PHQ8 = adapted version of Patient Health Questionnaire 9 (over 8 items); SF36-MCS = 36-item Short Form Health Survey, Mental Component Score.

**a)**

| Variables | INFLA-score | CRP | Plt | WBC | GLR |
| --- | --- | --- | --- | --- | --- |
| Socio-demographic COVs | 0.023  (0.011) | -0.007  (0.42) | 0.27  (0.61) | 0.004  (0.076) | 0.027  (< 10^-4^) |
| Socio-demographic COVs  +  Health Conditions MEDs | 0.022  (0.012) | -0.008  (0.35) | 0.40  (0.46) | 0.004  (0.098) | 0.027  (< 10^-4^) |
| Socio-demographic COVs  +  Lifestyle MEDs | 0.006  (0.45) | -0.015  (0.050) | 0.19  (0.73) | -0.002  (0.34) | 0.022  (7 x 10^-4^) |
| ALL COVs and MEDs | 0.006  (0.46) | -0.015  (0.048) | 0.32  (0.56) | -0.002  (0.27) | 0.021  (1.2 x 10^-3^) |

**b)**

| Biomarker | CPR | Plt | WBC | GLR | MPV | PCT | PDW | RDW | LY | MO | GR | NE | NLR |
| --- | --- | --- | --- | --- | --- | --- | --- | --- | --- | --- | --- | --- | --- |
| PHQ2 | -0.015  (0.048) | 0.32  (0.56) | -0.002  (0.27) | 0.021  (1.2 x 10^-3^) | -0.024  (0.019) | -0.0003  (0.54) | 0.036  (0.015) | 0.005  (0.65) | -0.195  (2.2 x 10^-3^) | -0.045  (0.013) | 0.239  (4 x 10-4) | 0.225  (5.6 x 10^-3^) | 0.023  (2.1 x 10^-3^) |

**Table S5.** Results of **a)** stepwise glm regressions of INFLA-score and it component biomarkers and **b)** full glm model regressions of different inflammation-related circulating biomarkers, vs Patient Health Questionnaire 2 (PHQ2; N = 13,492). ^a^ Beta (β) values referring to standardized psychometric scales and INFLA-score (but not standardized markers) are reported, along with association p-values in brackets. Abbreviations: COVs = covariates (confounders, see Table S1a above); MEDs = potential mediators tested; CRP = C-reactive protein (mg/L, log scale); Plt = platelet count (x10^9^/L); WBC = white blood cell count (x10^9^/L, log scale); GLR and NLR = granulocyte- and neutrophil-to-lymphocyte ratio; MPV = mean platelet volume (fL); PCT = plateletcrit (%); PDW = platelet distribution width (%); RDW = red cell distribution width (%); LY, MO, GR and NE = fraction of lymphocytes, monocytes, granulocytes and neutrophils over the total WBC (%).
